# The CATP-8/P5A-type ATPase functions in multiple pathways during neuronal patterning

**DOI:** 10.1101/2021.03.10.434729

**Authors:** Leo T.H. Tang, Meera Trivedi, Jenna Freund, Christopher J. Salazar, Nelson J. Ramirez-Suarez, Garrett Lee, Maisha Rahman, Yu Wang, Barth D. Grant, Hannes E. Bülow

## Abstract

The assembly of neuronal circuits involves the migrations of neurons from their place of birth to their final location in the nervous system, as well as the coordinated growth and patterning of axons and dendrites. In screens for genes required for patterning of the nervous system, we identified the *catp-8/P5A-ATPase* as an important regulator of neural patterning. P5A-ATPases are part of the P-type ATPases, a family of proteins known to serve a conserved function as transporters of ions, lipids and polyamines in unicellular eukaryotes, plants, and humans. While the function of many P-type ATPases is relatively well understood, the function of P5A-ATPases in metazoans remained elusive. We show here, that the *Caenorhabditis elegans* ortholog *catp-8/P5A-ATPase* is required for specific aspects of nervous system development. Specifically, the *catp-8/P5A-ATPase* serves functions in shaping the elaborately sculpted dendritic trees of somatosensory PVD neurons. Moreover, *catp-8/P5A-ATPase* is required for axonal guidance and repulsion at the midline, as well as embryonic and postembryonic neuronal migrations. Interestingly, not all axons at the midline require *catp-8/P5A-ATPase*, although the axons run in the same fascicles and navigate the same space. Similarly, not all neuronal migrations require *catp-8/P5A-ATPase*. A CATP*-8/P5A-ATPase* reporter is localized to the ER in most if not all tissues and *catp-8/P5A-ATPase* can function both cell-autonomously and non-autonomously to regulate neuronal development. Genetic analyses establish that *catp-8/P5A-ATPase* can function in multiple pathways, including the Menorin pathway, previously shown to control dendritic patterning in PVD, and Wnt signaling, which functions to control neuronal migrations. Lastly, we show that *catp-8/P5A-ATPase* is required for localizing select transmembrane proteins necessary for dendrite morphogenesis. Collectively, our studies suggest that *catp-8/P5A-ATPase* serves diverse, yet specific roles in different genetic pathways, and may be involved in the regulation or localization of transmembrane proteins to specific subcellular compartments.

**AUTHOR SUMMARY:** P-type ATPases are a large family of transporters that are conserved from unicellular eukaryotes and plants to metazoans. Structurally and functionally, they fall into five subfamilies, P1 to P5, of which the latter is further divided into P5A and P5B-type ATPases. Unlike for other P-type ATPases, no mutant phenotypes for the P5A-type ATPases have been described in metazoans. Here, we show that the *catp-8/P5A-ATPase* in the nematode *Caenorhabditis elegans* is involved in multiple aspects of neuronal patterning, including neuronal migrations as well as axon guidance and dendrite patterning. A functional fluorescent reporter fusion shows the CATP-8/P5A-ATPase is expressed in most, if not all, tissues in the endoplasmic reticulum and *catp-8* can function both in neurons and surrounding tissues from where it orchestrates neuronal development. Genetically, *catp-8* acts in multiple pathways during these processes, including the Wnt signaling and the Menorin pathway. Imaging studies suggest that the *catp-8/P5A-ATPase* is necessary for proper localization of cell-surface transmembrane molecules to dendrites of sensory neurons, but likely not for their trafficking. In summary, we propose that CATP-8/P5A-ATPase serves a function in the ER during development of select neurons, by localizing certain transmembrane, and possibly, secreted proteins

## INTRODUCTION

Development of a nervous system is a crucial step in metazoan development that requires the coordinated interactions of many genetic pathways and tissues. This process can be broken down into at least three components. First, in metazoans, a majority of neurons are actually born distant from their final location in adult animals. Therefore, neurons have to migrate from the place they are born to their final location in the nervous system, a process that is guided by both intrinsic and extrinsic factors [1, 2]. Second, axonal processes, the cellular structures that mediate output of nerve cells, need to grow and navigate, guided by extracellular guidance factors, towards their cellular targets with whom they form specific synaptic connections [3, 4]. Third, the often elaborately sculpted dendrites, a neuron’s receiving cellular structures, are patterned by dedicated pathways that originate from different tissues and act both cell-autonomously and non-autonomously [5–7].

The nematode is an excellent system to study basic aspects of conserved processes involved in neural patterning. Several cellular paradigms in worms can be studied at single cell resolution and often ample genetic information already exists for these processes. For example, the somatosensory PVD neurons are a powerful system to study conserved mechanisms of dendrite patterning [8, 9]. These mechanisms involve the interactions of different tissues that coordinately establish the stereotyped dendritic trees of PVD neurons. A complex of two transmembrane cell adhesion molecules, SAX-7/L1CAM and MNR-1/Menorin, function from the skin in concert with a secreted chemokine LECT-2/Chondromodulin II from muscle to instruct the growing PVD dendrites via a leucine rich transmembrane receptor DMA-1/LRR-TM expressed on the dendrites [8, 9]. Other well-studied paradigms include neuronal migrations, such as the anteriorly directed migration of hermaphrodite specific (HSN) neurons from the tail region towards the midbody region [10] or the pair of laterally located Q neuroblasts, which undergo stereotypical divisions and migrations during larval stages [11]. All cell migrations appear to be controlled, at least in part, by the conserved Wnt signaling system, among others [12, 13].

The diverse P-type family of transporters is a highly evolutionarily conserved family of transporters in eukaryotes that, respectively, move multiple molecular species, including ions and lipids across membranes through the hydrolysis of ATP. These P-type ATPases are multi-transmembrane proteins that can be divided into five subfamilies based on structural and functional characteristics, P1 through P5 [14, 15], of which the latter can be further divided into the P5A and P5B family [16]. The P1 through P3 subfamilies serve as ion transporters, whereas the P4 subfamily has been shown to function as flippases that move lipids across membranes [15]. The P5B family of transporters was recently shown to transport polyamines across membranes [17]. Mutations in the P5A family of transporters in both plants (*A. thaliana*) and unicellular eukaryotes (*S. cerevisiae*) were shown to display pleiotropic phenotypes, including defects in phosphate homeostasis and male gametogenesis [18–20], as well as defects in phospholipid and sterol homeostasis [21] and targeting of mitochondrial outer membrane proteins [22], respectively. However, it remained unclear what functions P5A-ATPases serve in metazoans.

We show here that the sole *C. elegans* ortholog *catp-8/P5A-ATPase* is required for specific aspects of nervous system development, including in dendrite patterning, axonal guidance decisions and neuronal migrations. A CATP-8/P5A-ATPase reporter shows expression in the endoplasmic reticulum in most if not all tissues albeit at different levels. Moreover, the *catp-8/P5A-ATPase* can function both cell-autonomously and non-autonomously in multiple genetic pathways in different cellular contexts. Additionally, we find that in *catp-8/ATPase* mutants, the amounts of functional reporters for select transmembrane proteins (the leucine rich transmembrane receptor DMA-1/LRR-TM and the HPO-30/Claudin-like molecule) is strongly reduced on higher order dendritic branches of PVD dendrites. Taken together, these results suggest that the CATP-8/P5A-ATPase functions in multiple pathways during neural patterning and is important for correct localization of transmembrane cell surface proteins. While this study was in progress, two studies reporting a function for *catp-8/P5A-ATPase* in PVD patterning in *C. elegans* were published [23, 24]. Feng et al. suggested a role for *catp-8/P5A-ATPase* in targeting of DMA-1/LRR-TM in dendrites [23] and Qin et al. suggested that *catp-8* was important for ER homeostasis by preventing mistargeting of mitochondrial proteins independently of ER-associated degradation (ERAD) [24]. Similarly, a crystal structure of the yeast homolog Spf1 in conjunction with biochemical studies suggested a role for P5A-ATPases in ER and mitochondrial quality control [25].

## RESULTS

### The P5A-type ATPase CATP-8 is required for somatosensory dendrite patterning

In a genetic screen for animals with defects in dendrite arborization of PVD somatosensory neurons (Ramirez-Suarez *et al.*, unpublished), we isolated the allele *dz224* with characteristic defects in dendrite patterning (see below). Using a combination of whole genome sequencing and mapping (Fig.S1, Materials and Methods)[26]), we found that the molecular lesion in *dz224* results in a premature stop codon after 55 amino acids in *catp-8*, which encodes the sole P5A-type ATPase in the *C. elegans* genome (Fig.1A-C). Based on the molecular nature of *dz224*, we predict this allele to result in strong, if not complete loss of function of the *catp-8/P5A-ATPase*.

**Figure 1.**
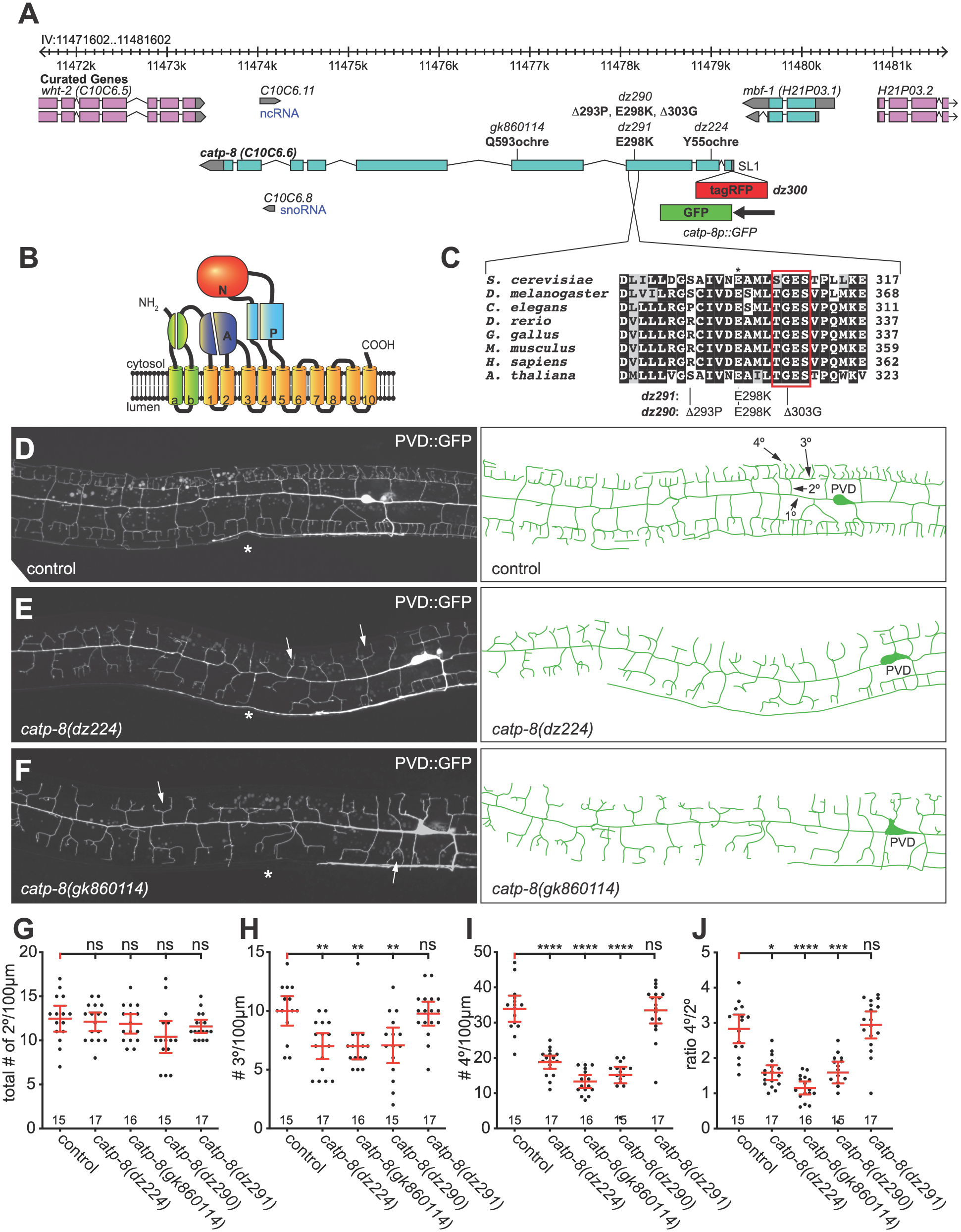
CATP-8/P5A-ATPase is required for PVD dendrite patterning. A. Genetic environs of the *catp-8* locus on chromosome IV. *Catp-8* is encoded on the minus strand and transcribed from right to left. Alleles used in this study are indicated as are the resulting molecular changes. The *catp-8* transcript can be SL1 spliced based on the *yk1435b05* EST clone (gift of Y. Kohara), suggesting that *catp-8* may be part of a hybrid operon. A schematic of the *catp-8p::GFP* transcriptional reporter is shown as is the genomic location of the tagRFP insertion to create the translational tagRFP::CATP-8 reporter. B. Schematic domain structure and membrane topology of the CATP-8/P5A ATPase with the following domains indicated: A: actuator domain, N: nucleotide binding domain, P: phosphorylation domain. C. Alignment of a conserved section of the actuator domain, which encodes a functional phosphatase function that is required for functional cycling of P-type ATPases [15]. The E298K mutation (asterisk, equivalent to the E349K mutant associated with intellectual disability [32]) is immediately adjacent to the (S/T)GES loop (boxed in red), which is important for A-domain function [25]. Noted on the right is the number of amino acids. The TGES motif is boxed in red. Accession numbers of sequences for P5A-type ATPases used in the alignment are P39986 (*S. cerevisiae*), Q9VKJ6 (*D. melanogaster*), P90747 (*C. elegans*), F1R1X4 (*D. rerio*), Q5ZKB7 (*G. gallus*), Q9EPE9 (*M. musculus*), Q9VKJ6 (*H. sapiens*), Q9LT02 (*A. thaliana*). D. -F. Fluorescent micrographs of PVD neurons with corresponding schematics on the right in the genotypes indicated. PVD was visualized with *wdIs52 Is[F49H12.4p::GFP]* [28]. Asterisks mark the location of the vulva. In D., primary (1°), secondary (2°), tertiary (3°) and, quaternary (4°) dendrites are indicated. White arrows in E. and F. indicate typical *catp-8* mutant defects. G. - J. Quantification of the number of secondary (G), tertiary (H), and quaternary (I) dendrite branches, as well as the ratio of quaternary to secondary (J) branches, 100µm anterior to the PVD cell body in the genotypes indicated. Data are represented as the mean ± 95% confidence interval. Statistical significance was calculated using one-sided ANOVA with Tukey’s multiple comparison test. *, *P* ≤ 0.05; **, *P* ≤ 0.005; ***, *P* ≤ 0.0005; ****, *P* ≤ 0.0001; ns, not significant. n = 15 wild-type control (*wdIs52*) animals; n = 17 *catp-8(dz224)* mutant animals; n = 16 *catp-8(gk860114)*; n = 15 *catp-8(dz290)*; n = 17 *catp-8(dz291)*.

To more precisely determine the function of *catp-8* in dendrite patterning, we conducted morphometric analyses of PVD somatosensory dendrites. During larval stages two primary dendrites emanate from the PVD cell body on either side of the animal in an anterior and posterior direction, respectively, and through sequential perpendicular branching of secondary, tertiary, and quaternary branches establish highly stereotyped dendritic trees with semblance to Menorah-like candelabras (Fig.1D)[27–29]. We traced dendrites in a region 100µm anterior to the cell body of young adult animals and determined the number and length of secondary, tertiary and quaternary dendritic branches. Compared to wild type animals, we found the number of tertiary and quaternary branches to be significantly reduced in *catp-8(dz224)* mutants (Fig.1D-J), while the total number of secondary branches remained unchanged. However, the number of secondary branches that reached the line of the tertiary branches was significantly reduced (Fig.S1B). Overall, many dendritic trees displayed a conspicuous defect, where secondary branches appeared to lack proper tertiary branches, and merely bifurcated into two quaternary dendrites rather than forming perpendicular tertiary branches. This observation was also borne out by a reduced number of quaternary branches per secondary branches, approaching an average of two quaternary branches per secondary branch (i.e. per menorah) as opposed to approximately four in control animals and, a decrease in aggregate length of secondary, tertiary and quaternary branches (Fig.1K, Fig.S1B-E). In addition, we observed increased numbers of ectopic secondary and ectopic tertiary branches, possibly reflective of a lack of a functioning feed-back loop that has been postulated to restrict secondary branching once stable tertiary branches have been formed [30](data not shown).

To corroborate these findings, we obtained three additional alleles of *catp-8/P5A ATPase*, including *gk1860114* from the million mutation project [31], which results in a premature stop codon after 593 amino acids (Fig.1A). We used CRISPR/Cas9 based genome editing to create two additional alleles (Fig.1A): *dz290*, which deletes two codons within a perfectly conserved stretch of the actuator domain [15], and a missense mutation (E289K), which changes a perfectly conserved glutamic acid to lysine (E289K) in the actuator domain (Fig.1A), and is analogous to a mutation described in a pair of sisters with intellectual disability [32]. Finally, we introduced the E289K mutation on its own to create *dz291*. Of these, all but the *dz291* allele displayed strong phenotypes in PVD patterning that were indistinguishable from the *dz224* allele (Fig.1G-K, Fig.S1B-E). Consistent with the nonsense mutations in *dz224* and *gk860114*, both alleles behaved recessively and failed to complement each other, suggesting loss of *catp-8* function (Fig.S1F). To test, whether the *gk860114* allele represented the null phenotype, we analyzed animals where *catp-8(gk860114)* was placed in trans to the *mDf7* deficiency, which removes the genomic region encoding *catp-8* and fails to complement *catp-8* mutant alleles (Fig.S1A). We found that these animals did not display a more severe phenotype in PVD patterning than *catp-8(gk860114)* homozygous animals, demonstrating that *gk860114* behaves as a genetic null allele (Fig.S1H). Animals carrying the *catp-8(gk860114)* null allele appear superficially normal, have a normal body length at the L1 larval stage (Fig.S1I), but seem to grow slower (data not shown). Taken together these findings suggest that *catp-8* function is required for PVD patterning, and especially for the extension of tertiary and quaternary dendritic branches. Moreover, the E289K mutation, which is analogous to a mutation shared by the pair of sisters with intellectual disability [32], does not compromise *catp-8* sufficiently to result in morphological phenotypes, at least in the context of PVD patterning or neuronal migration (see below, data not shown). Lastly, the defects due to mutations in the actuator domain are consistent with the interpretation that enzymatic activity of CATP-8/P5A type ATPase is required for PVD patterning.

### The P5A-type ATPase CATP-8 is required for specific neuronal and axonal migrations

We next asked whether *catp-8* is only necessary for development of the highly branched dendritic trees of PVD dendrites or, more generally for other aspects of neural patterning. To this end, we crossed *catp-8* mutants with a panel of GFP reporters that label individual neurons or classes of neurons, including motor neurons, interneurons, sensory neurons and touch receptor neurons. We found that mutant alleles of *catp-8* displayed defects in neurite patterning of some but not all neurons. For example, in *catp-8* mutant animals the D-type motor neurons displayed neurite extension defects in the ventral and dorsal nerve cords, whereas the DA/DB-motor neurons appeared indistinguishable from control animals (Table 1, Fig.S2A-D). This is significant insofar as the processes of both types of motor neurons make similar navigational choices and travel, at least in the nerve cords in the same fascicles. However, neither patterning of the pair of AIY interneurons, nor the amphid sensory neurons or phasmid sensory neurons was obviously dependent on *catp-8* function (Table 1, Fig.S2E-J). Lastly, we investigated axon guidance choices of HSN and PVQ neurons at the ventral midline, where both neurons send axons in the left and right ventral nerve cords from the midbody or tail to the head region, respectively. We observed significant defects in axon midline guidance of HSN neurons with frequent cross overs, but no defects for PVQ neurons (Table 1, Fig.S2K-N). Taken together, our findings show that axonal patterning of some but not all neurons is dependent on *catp-8/P5A-ATPase* function. Importantly, even neurons that make similar guidance choices and travel in the same nerve fascicles such as the D-type and DA/DB motor neurons or the HSN and PVQ neurons, can develop either dependent on *catp-8* function or, independently.

**Table 1.** Survey of neuroanatomical defects in catp-8/P5A-ATPase mutant animals.

| **Neurons Examined (Marker Used)** | **wild type**<br>**(% defect)** | **N** | ***catp-8(0)***<br>**(% defect)** | **N** | **P Value<sup>1</sup>** |
| --- | --- | --- | --- | --- | --- |
| **Motor Neurons<sup>2</sup>** |  |  |  |  |  |
| <u>D-Type (*juls76*)</u> |  |  |  |  |  |
| Gaps in ventral nerve cord | 10.8 | 93 | 41.3 | 109 | **** |
| Gaps in dorsal nerve cord | 12.9 | 93 | 26.6 | 109 | * |
| L/R choice of commissures | 35.6 | 115 | 50 | 144 | * |
| <u>DA/DB (*evls82b*)</u> |  |  |  |  |  |
| Gaps in ventral nerve cord | 0 | 102 | 0 | 108 | ns |
| Gaps in dorsal nerve cord | 0 | 102 | 0 | 108 | ns |
| L/R choice of commissures | 4.8 | 102 | 5.7 | 108 | ns |
| <u>HSN (*dzls75*)</u> |  |  |  |  |  |
| Ventral nerve cord crossover | 28.4 | 64 | 48.3 | 118 | ** |
| **Interneurons<sup>2</sup>** |  |  |  |  |  |
| <u>PVQ (*oyls14*)</u> |  |  |  |  |  |
| Ventral cord crossover | 11.8 | 101 | 7 | 100 | ns |
| <u>AIY (*mgls32*)</u> |  |  |  |  |  |
| Early termination | 0 | 120 | 0 | 124 | ns |
| Ectopic axonal branching | 2 | 120 | 2 | 124 | ns |
| **Amphid Neurons<sup>3</sup>** |  |  |  |  |  |
| <u>Dil Stain</u> |  |  |  |  |  |
| Defasciculation | 6 | 50 | 4 | 50 | ns |
| Early termination | 0 | 50 | 0 | 50 | ns |
| Anterior process | 0 | 50 | 0 | 50 | ns |
| Anterior nerve ring | 0 | 50 | 0 | 50 | ns |
| Posterior process | 0 | 50 | 0 | 50 | ns |
| Posterior nerve ring | 0 | 50 | 0 | 50 | ns |
| **Phasmid Neurons<sup>3</sup>** |  |  |  |  |  |
| <u>Dil Stain</u> |  |  |  |  |  |
| Ectopic axons | 0 | 50 | 0 | 50 | ns |
| Axon crossover | 0 | 50 | 0 | 50 | ns |
| Wandering processes | 0 | 50 | 0 | 50 | ns |

<sup>1</sup> Statistical significance was calculated using . \*, $P \leq 0.05$ ; \*\*, $P \leq 0.005$ ; \*\*\*\*, $P \leq 0.0001$ ; ns, not significant.
<sup>2</sup> Neuroanatomical defects were scored as described [63]
<sup>3</sup> Neuroanatomical defects were scored as described [64]

We next investigated a set of stereotyped neuronal and non-neuronal cellular migrations, including both embryonic and postembryonic migrations, as well as anteriorly-posteriorly and posteriorly-anteriorly directed migrations (Fig.2A). We first investigated two embryonic migrations: the pair of ALM mechanosensory neurons, which migrate from the head region towards the midbody region (i.e. anteriorly-posteriorly directed) and, the hermaphrodite specific neurons (HSNs), a set of motor neurons that migrate from the tail towards the mid body region (i.e. posteriorly-anteriorly directed)[33]. Loss of *catp-8* function resulted in severe migration defects in anteriorly directed HSN migrations, which often failed to reach their final destination around the midbody and, instead stopped migration prematurely (Fig.2B,E). In contrast, loss of *catp-8* had no effect on posteriorly directed ALM neuronal migrations (Fig.2C,F). We next investigated a set of postembryonic migrations, specifically of the Q cell descendants [11]. The bilateral pair of Q cells are neuroblast cells, which are born during the mid L1 larval stage. On the right side of the animal, the QR cell migrates anteriorly while undergoing a series of stereotyped cell divisions as follows. The first division results in two more blast cells, of which one divides again and gives rise to the AQR cell and the other undergoes programmed cell death [34]. The other blast cell also gives rise to one cell that undergoes programmed cell death and another that after a further division forms the AVM and SDQR neurons (Fig.2A). The AQR or AVM and SDQR neurons therefore originate from two subbranches of QR cell descendants, respectively, and are positioned in the head of the animals or in a lateral anterior section of the worms, respectively. The Q cell descendants on the left side of the animal undergo a posteriorly directed migration, while also giving rise, in a similar pattern of divisions, to three neurons, PQR or PVM and SDQL (Fig.2A). PQR travels to the tail whereas PVM and SDQL remain in a lateral posterior section of the animals. We found that the anteriorly directed migrations of both AQR and AVM, which represent both sub branches of QR descendants, are not obviously affected in *catp-8* mutants, whereas the posteriorly directed migrations of PQR and PVM neurons display defects (Fig.2C,D,G,H). Specifically, PQR and PVM erroneously migrate anteriorly towards the head instead of posteriorly. This observation suggests that the choice of directionality in migration of QL descendants on the left side of the animals is dependent on *catp-8/P5A ATPase* function, rather than migratory capacity itself. Moreover, we found that *gk860114* homozygous mutants originating from homozygous mothers displayed Q cell migration defects, whereas *gk860114* homozygous mutants originating from *gk860114/+* heterozygous animals did not display this phenotype, suggesting a maternal contribution of *catp-8* during this postembryonic migration (Fig.S1G). Lastly, we investigated two pairs of non-neuronal cells, the coelomocytes, which migrate embryonically in a posterior direction [33]. We found that coelomocyte migration proceeds normally in *catp-8* mutants (Fig.2I). Collectively, our experiments show that both embryonic and postembryonic cell migrations require *catp-8/P5A-ATPase* function, including anteriorly and posteriorly directed migrations.

**Figure 2.**
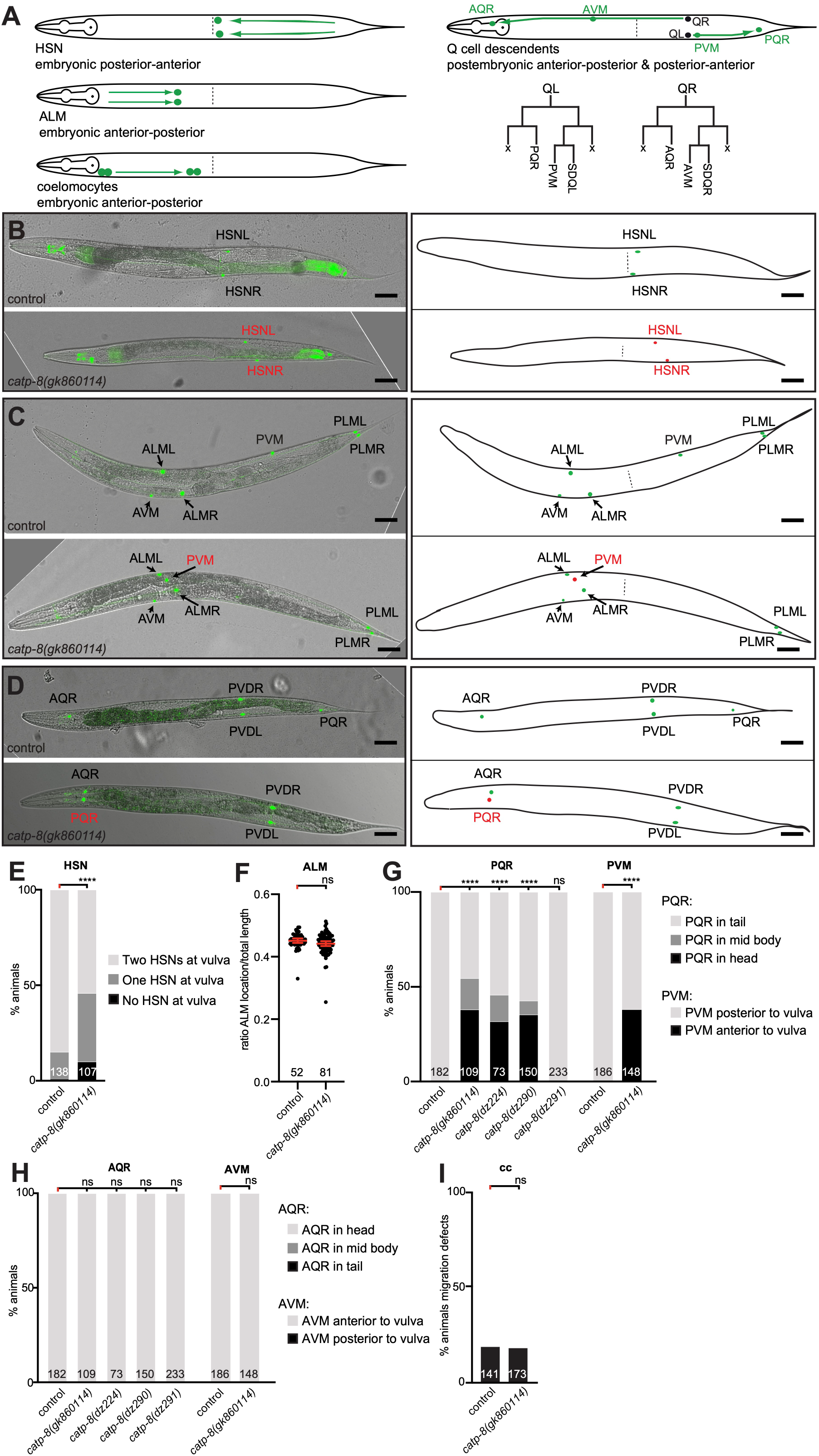
CATP-8/P5A-ATPase is required for neuronal migrations. A. Schematics of different cellular migrations. Shown on the left are embryonic migrations including HSN (posterior to anterior), ALM (anterior to posterior) and coelomocytes (anterior to posterior). For simplicity, migrations are illustrated in adult animals. Indicated on the right are the migrations of the descendants of the QL and QR neuroblasts, respectively, which arise in stereotyped cell-divisions as indicated below. Note that Q cells divide into distinct sub-branches, comprising the AQR/PQR or AVM/PVM and SDQR/SDQL neurons, respectively. X denotes cells that undergo programmed cell death [34]. B. - D. Composite of fluorescent and differential interference contrast images of reporters visualizing the HSN neurons (B), touch receptor neurons (C) or AQR/PQR (D). Images of wild type and *catp-8(gk860114)* mutant animals are shown in top and lower panels, respectively, with corresponding schematics on the right. Correctly positioned neurons are shown in green and incorrectly positioned neurons in red. HSN visualized by *dzIs75 II Is[kal-1p9::GFP; ttx-3p::mCherry]*[40]), AQR/PQR by *wdIs52 II Is[F49H12.4p::GFP]*, and touch receptor neurons by *zdIs5 I Is[mec-4p::GFP]* [65]. Scale bar = 25µm. E. Quantification of cell migration defects of HSN neurons. Percent animals are shown with neurons in wild type position (light grey bars), partially incomplete migration (dark grey bars) and completely failed migration (black bars). Pairwise statistical significance was calculated using Fisher’s exact test. **** *P* ≤ 0.0001; ns, not significant. F. Quantification of ALM migration comparing ratio of ALM location to total animal length in control versus *catp-8(gk860114)*. Unpaired t test, ns, not significant. G. - H. Quantification of migration of PQR and PVM (G) and AQR and AVM (H) neurons comparing control versus indicated alleles of *catp-8*. Percent animals are shown with neurons in wild type position (light grey bars), partially incomplete migration (dark grey bars) and completely failed migration (black bars). Pairwise statistical significance was calculated using Chi-squared test or Fisher exact test. **** *P* ≤ 0.0001; ns, not significant. I. Percentage of animals exhibiting coelomocyte migratory defect in wildtype or *catp-8(gk860114)*. Fisher’s exact test. ns not significant.

### CATP-8/P5A ATPase functions both cell-autonomously and non-autonomously in neural patterning

To gain insight into the cellular focus of action of the *catp-8/P5A-ATPase*, we used CRISPR/Cas9 engineering to insert a tagRFP fluorescent protein at the N-terminus of CATP-8. We found these animals to show widespread, if not ubiquitous expression of CATP-8, including in the epidermis, muscle, pharynx, intestine, and the major neuronal ganglia (Fig.3A-D). The expression appears intracellular, in a perinuclear pattern and, different tissues appear to express different levels of CATP-8. Of note, we detected expression in both neurons that were phenotypically affected in *catp-8* mutants as well as neurons that were not visibly affected. For example, clear expression of the endogenous tagRFP::CATP-8 reporter was seen in HSN, PVM, and PVD neurons, but also in ALM neurons and others (Fig.3E-H, data not shown). Importantly, PVD morphology and PQR cell position appeared completely normal in animals carrying the endogenous fluorescent tag, suggesting that the N-terminal tagRFP::CATP-8 fusion is fully functional. We found similar expression patterns in transgenic animals expressing GFP under control of the *catp-8* 5’ region immediately upstream of the ATG start codon, where we also saw widespread expression from embryonic stages on, including in muscle, intestine, pharynx and other tissues (Fig.S3).

**Figure 3.**
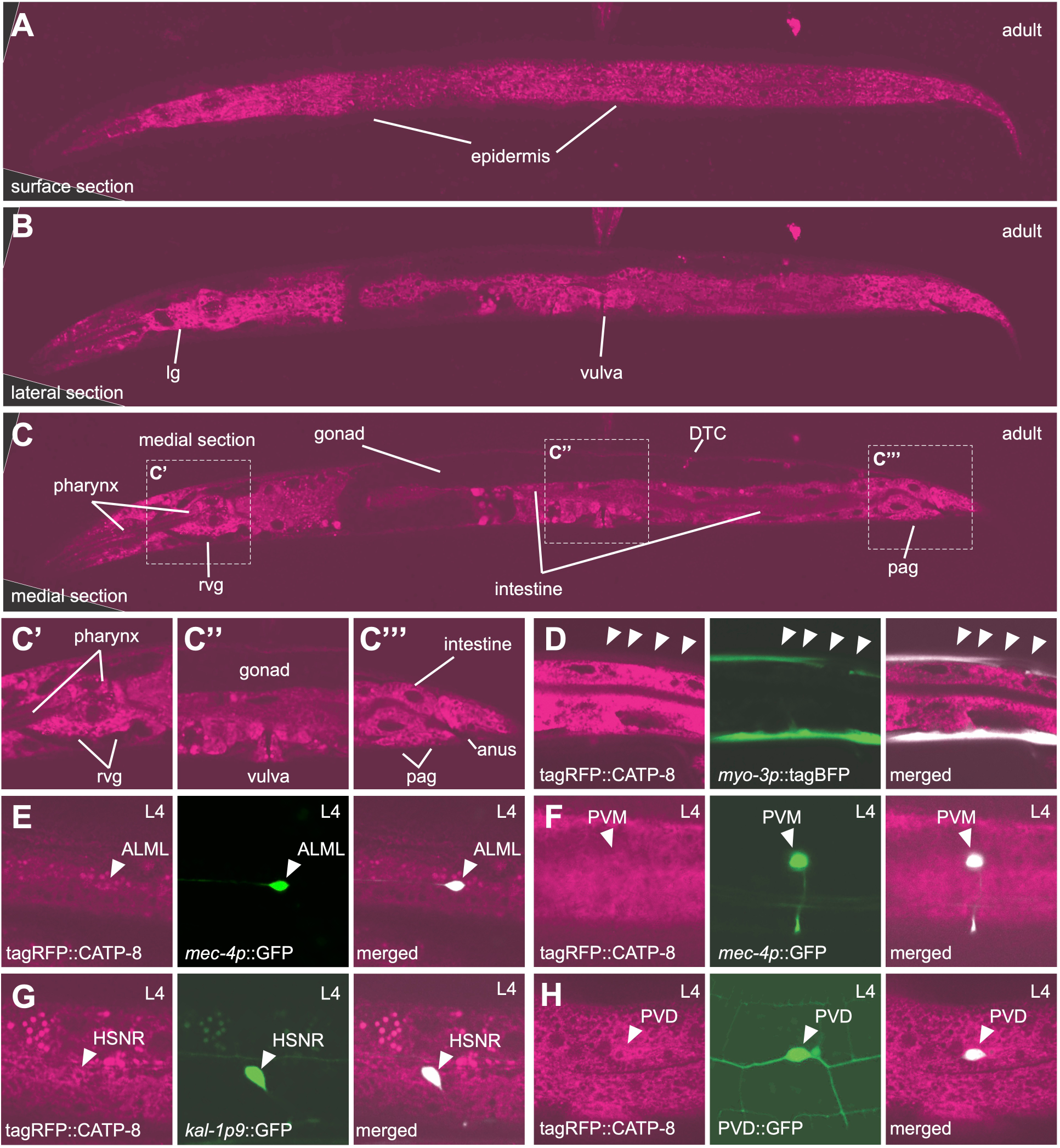
CATP-8/P5A-ATPase is widely expressed at apparently different levels. A.- C. Whole body confocal optical sections of tagRFP translational knock-in of *catp-8* animals, showing perinuclear expression in tissue indicated. An optical section near the surface of the animals shows epidermal expression, whereas a more lateral and a medial section reveal many other tissues, including pharynx, intestine, lateral ganglion (lg), retrovesicular ganglion (rvg), distal tip cell (DTC), and preanal ganglion (pag). Insets in C are shown magnified as C’, C’’, C’’’. Note that tagRFP::CATP-8 is not visibly expressed in the gonad and that expression levels vary in different tissues (e.g. compare epidermal and muscle expression, confer with D). D.- H. Confocal images showing CATP-8 expression in indicated neurons and tissues, as evident by colocalization of tagRFP::CATP-8 with relevant markers, including *dzEx2136* (*Ex[myo-3p::tagBFP; mec-4p::catp-8]* from muscle, D), *zdIs5* (*Is[mec-4p::GFP]* for ALM and PVM, E, F), *dzIs75* (*Is[kal-1p9::GFP]* for HSN, G), *wdIs52* (*Is[F49H12.4p::GFP]* for PVD, H).

To determine, where *catp-8* may function during different developmental processes, we performed transgenic rescue experiments, where we expressed a *catp-8* cDNA under control of heterologous promoters in *catp-8* mutant animals. We found that patterning defects of PVD dendrites were rescued by pan-neuronal transgenic expression of *catp-8*, but not by expression in muscle (Fig.4A-G, Fig.S4A-C). Interestingly, a single copy transgene expressing the functional N-terminal tagRFP::CATP-8 fusion under control of the heterologous epidermis-specific *dpy-7p* promoter also rescued PVD patterning defects in *catp-8* mutants (Fig.4A-G, Fig.S4A-C). These observations suggest that *catp-8* can serve redundant functions in different tissues, i.e. both cell-autonomously and non-autonomously, but may be sufficient in either neurons or the epidermis. Surprisingly, embryonic and postembryonic cell migration defects of HSN, PQR and, PVM neurons were completely rescued by expression of *catp-8* in muscle, but not in the epidermis, the touch receptor neurons (i.e. PVM), or the precursors that give rise to the Q cells (Fig.4H-J, Fig.S4D-E). One out of three transgenic lines expressing the *catp-8* cDNA under control of a pan-neuronal reporter showed partial rescue of the neuronal migration defects of PVM and PQR neurons (Fig.4H-J, Fig.S4E), suggesting that overexpression in neurons can partially compensate for loss of *catp-8* in other tissues, e.g. muscle. We conclude that at least for these cellular migrations, *catp-8* appears to function non-autonomously in muscle. Collectively, our studies show that *catp-8* is widely if not ubiquitously expressed, albeit at different levels. Furthermore, *catp-8* can serve both autonomous and non-autonomous functions during neural patterning, including in dendrite morphogenesis, axonal guidance and different neuronal migrations.

**Figure 4.**
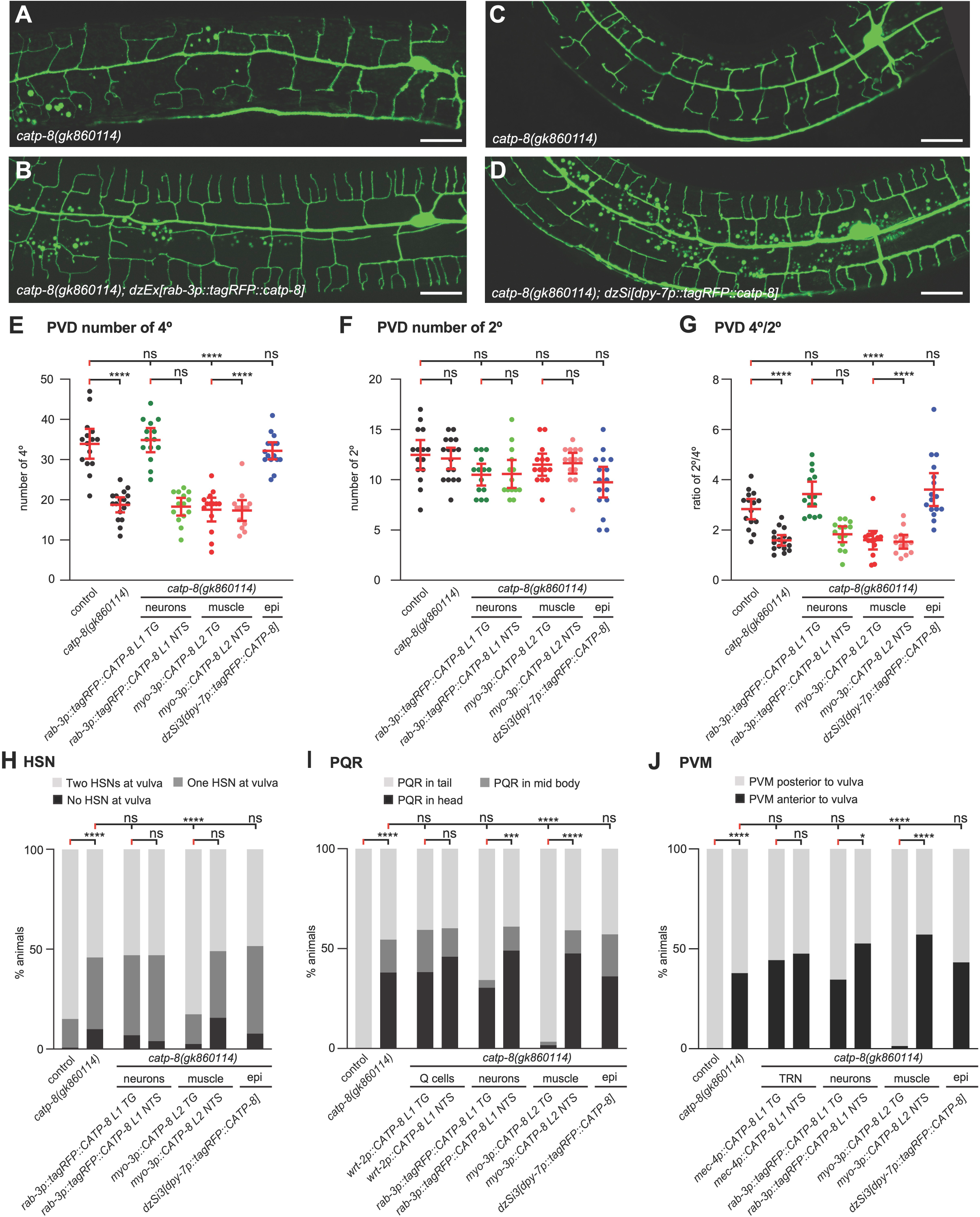
CATP-8/P5A-ATPase can function cell autonomously and non-autonomously during neural patterning. A. - B. Maximum z-projection of confocal images of PVD::GFP marker *wdIs52* from non-transgenic (A) and transgenic (B) sibling animals carrying pan-neuronal *catp-8* rescue array under *catp-8* mutant background, showing restoration of PVD structure under neuronal rescue. Scale bar = 20µm. C. - D. Maximum z-projection of confocal images of PVD::GFP marker *wdIs52* from animals without (C) and with (D) single-copy insertion of epidermal *catp-8* rescue under *catp-8* mutant background, showing restoration of PVD structure under epidermal rescue. Scale bar = 20µm. E. - G. Quantification of number of quaternary branches (E), secondary branches (F), and the ratio of quaternary to secondary branches (G) 100µm anterior to the PVD cell body in animals of genotypes indicated, showing complete rescue of *catp-8(gk860114)* phenotype when *catp-8* is expressed from pan-neuronal or epidermal promoter, but not muscle promoter. Data are represented as the mean ± 95% confidence interval. **** P<0.0001, ns, not significant, Kruskal-Wallis test with Dunn’s multiple comparisons test. n = 14 animals per genotype. H. - J. Percentage of animals with the indicated migration phenotype of HSN (D), PQR (E) and PVM (F) under indicated genotype backgrounds, showing all migration phenotypes of *catp-8* were rescued completely when *catp-8* is expressed from the muscle, and fail to be rescued when expressed cell-autonomously, pan-neuronally or epidermally. **** *P < 0.0001*, ns not significant, Chi-squared test. n > 75 animals per genotype.

**Figure 5.**
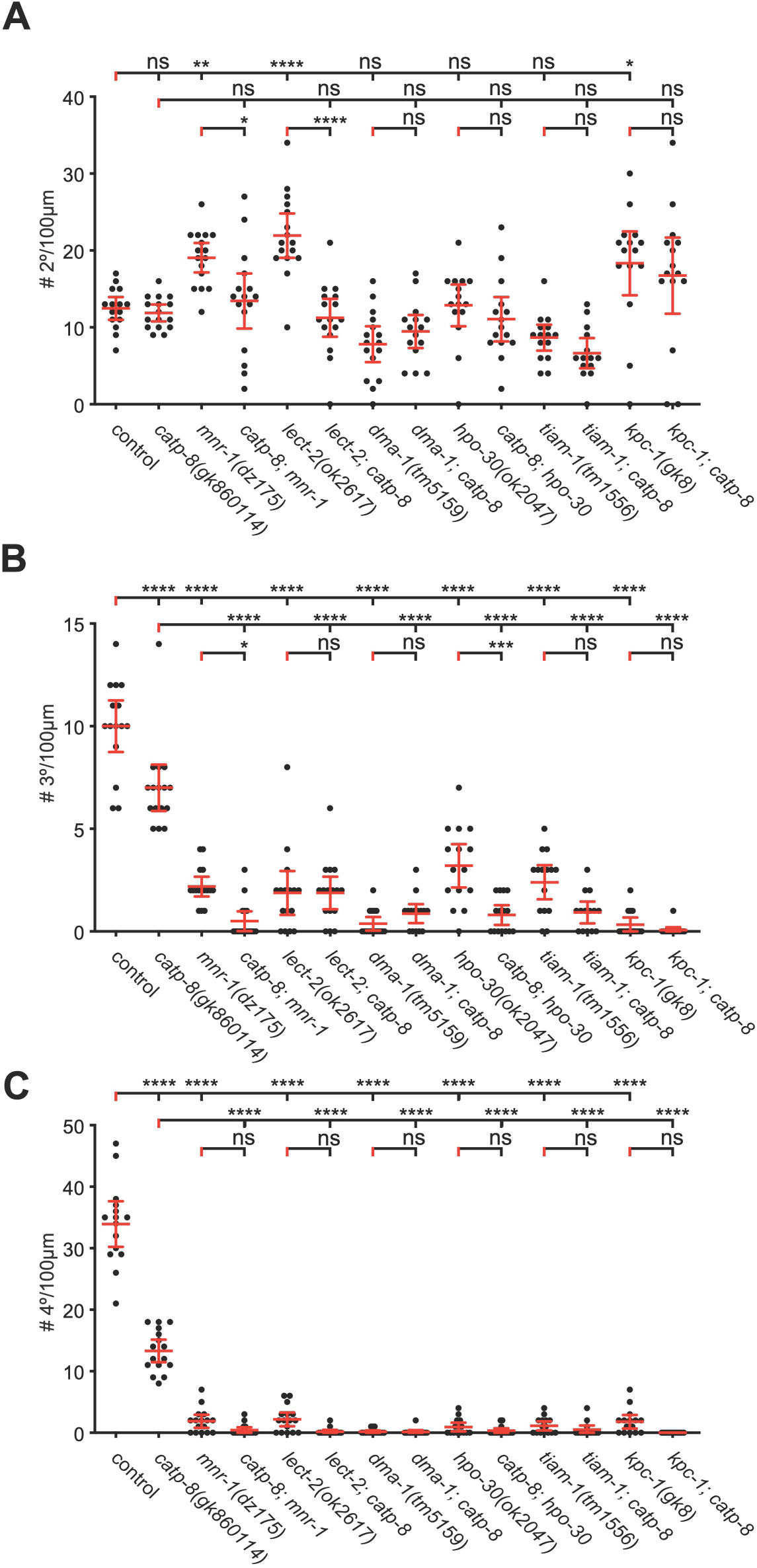
Genetic interactions of *catp-8/P5A ATPase* with the menorin pathway during dendrite patterning. Quantification of the number of secondary (A), tertiary (B), and quaternary (C) dendrite branches within 100µm anterior to the PVD cell body in animals of the indicated genotypes. Note that all alleles are molecular or genetic null alleles. Data are represented as the mean ± 95% confidence interval. *\* P<0.05, ** P<0.01, *** P<0.001, **** P < 0.0001* ns not significant; one-sided ANOVA with Tukey’s multiple comparison test. n = 15 animals per genotype.

### CATP-8/P5A ATPase functions in the Menorin pathway to pattern PVD dendrites

The conserved Menorin pathway shapes somatosensory patterning of PVD dendrites [8, 9]. Briefly, the two putative cell adhesion molecules SAX-7/L1CAM and MNR-1/Menorin form a complex in the epidermis, which, together with the muscle derived chemokine LECT-2/Chondromodulin II forms a high affinity substrate for the leucine rich repeat transmembrane receptor DMA-1/LRR-TM, which functions in PVD neurons. Downstream of DMA-1/LRR-TM in PVD function is a conserved set of intracellular molecules such as the TIAM-1/GEF guanine nucleotide exchange factor and HPO-30/Claudin, which together regulate the actin cytoskeleton [8, 35]. Previous genetic experiments established that SAX-7/L1CAM, LECT-2/Chondromodulin II and MNR-1/Menorin function in a linear genetic pathway with the DMA-1/LRR-TM receptor, but also suggested that DMA-1 can serve functions in PVD patterning that are independent of SAX-7/L1CAM, MNR-1/Menorin and LECT-2/Chondromodulin. To determine the genetic relationship between *catp-8* and the Menorin pathway, we analyzed double null mutants of *catp-8* and various components of the Menorin pathway using morphometric analyses. We traced and quantified the number and length of secondary, tertiary and quaternary dendrites 100µm anterior to the PVD cell bodies as previously described [30]. With regard to the number of secondary dendrites, we found that the *catp-8* mutant was epistatic in double mutants between *catp-8* and *mnr-1/Menorin* and *lect-2/Chondromodulin II*, i.e. the double mutant resembled the *catp-8* single mutant rather than displaying the increased number of secondary branches observed in the *mnr-1/Menorin* and *lect-2/Chondromodulin II* single mutants. In double mutants with *dma-1*, *hpo-30* or *tiam-1* the phenotypes were not further enhanced by concomitant genetic removal of *catp-8.* With regard to the number of tertiary branches, we made similar observations, although in some cases the double mutants between *catp-8* and for instance *mnr-1/Menorin*, or hpo*-30/Claudin* appeared phenotypically stronger than the single mutants (Fig.4B). For the number of quaternary branches, double mutants between *catp-8* and genes in the Menorin pathway invariably displayed the stronger phenotype observed in mutants of the Menorin pathway (Fig.4C). We observed similar genetic interactions when investigating the aggregate lengths of secondary, tertiary and quaternary branches (Fig.S5). We conclude that *catp-8* functions in the Menorin pathway and likely within both branches, i.e. the *mnr-1/Menorin-*dependent as well as -independent branches, both of which feed into the DMA-1/LRR-TM receptor in PVD neurons.

### CATP-8/P5A ATPase functions with the Wnt signaling pathway to control neuronal migrations

We found that *catp-8* is required for both embryonic and postembryonic neuronal migrations, including the anteriorly directed embryonic migrations of HSN neurons and the posteriorly directed postembryonic migrations of the Q cell descendants PVM and PQR. Both of these cell migrations are genetically well-described and depend on, among other pathways, Wnt signaling [10, 11, 13]. Briefly, the *egl-20/Wnt* ligand is expressed in posterior tissues, including body wall muscle [36, 37], believed to form a gradient [38] and function through the G-protein coupled receptor *mig-1/Frizzled* in HSN neurons to mediate HSN migration [39]. While other Wnts frizzled receptors play smaller, largely redundant roles in HSN migration [13], most functions appear to be mediated by the MIG-1/Frizzled receptor in HSN neurons, because a *mig-1; egl-20* double mutant does not display more severe defects in HSN migration than the single mutants [40]. To investigate the genetic relationship between *catp-8/P5A-ATPase* and Wnt signaling, we therefore created a double null mutant between *catp-8* and *mig-1.* We found that the HSN migration phenotype in *mig-1; catp-8* double mutant animals was not more severe than in either single mutant alone (Fig.6A), suggesting the *catp-8* functions genetically in a pathway with *mig-1/Frizzled* and, by inference, Wnt signaling.

**Figure 6.**
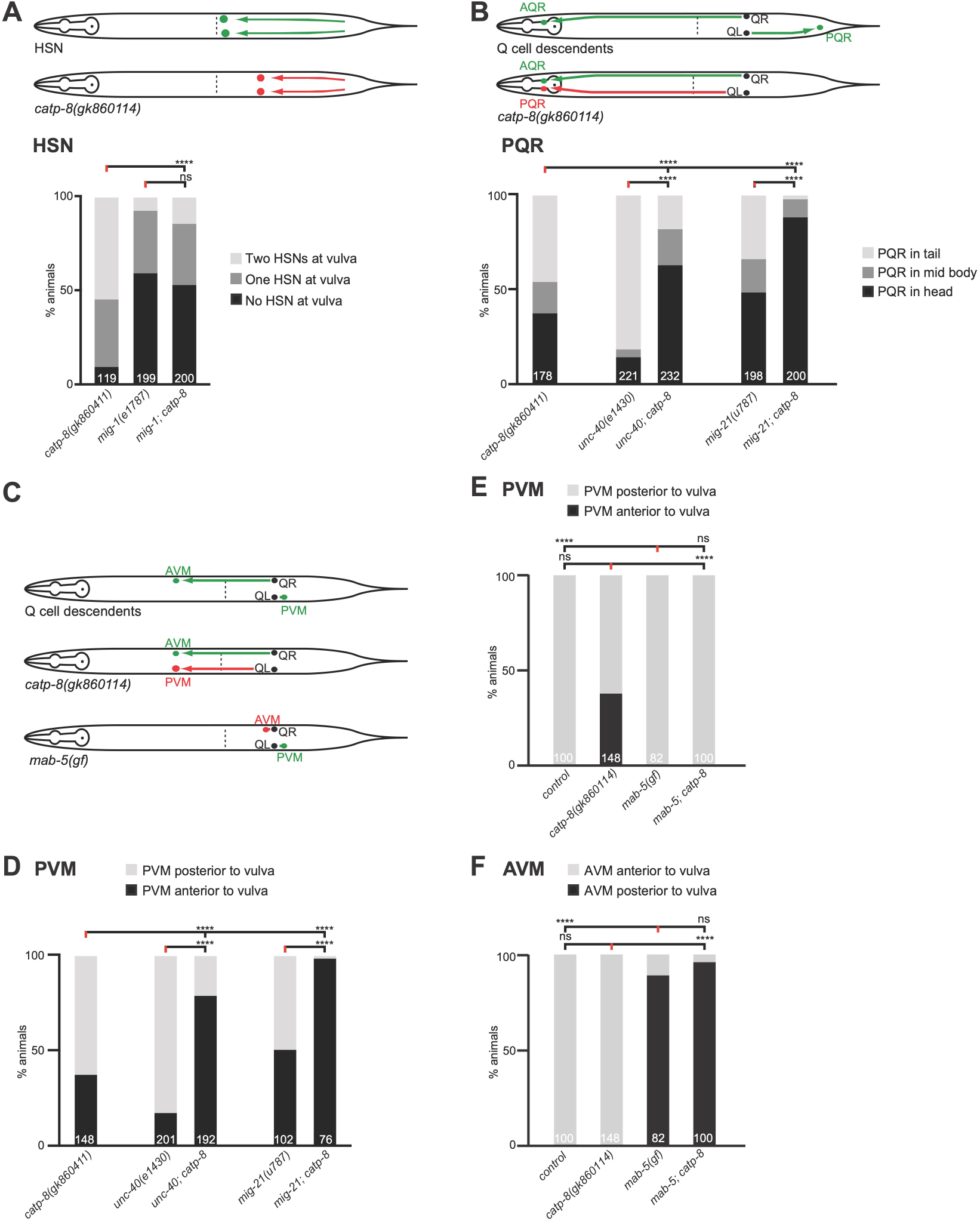
Genetic interactions of *catp-8/P5A* ATPase during cell migrations. A-C. Schematics depicting HSN (A), AQR/PQR (B), and AVM/PVM (C) migration (upper) and migration defects in indicated genotypes. A, B, D-F. Quantification of migration of HSN (A), AQR/PQR (B), PVM (D, E) and AVM (F) migration defects in indicated genotypes. Percent animals are shown with neurons in the wild type position (light grey bars), partially incomplete migration (dark grey bars) and completely failed migration (black bars). Pairwise statistical significance was calculated using Chi-squared test or Fisher exact test. ****, *P* ≤ 0.0001; ns, not significant.

The migration of Q cell descendants occurs in two phases [11]. In an initial phase, the QL and QR cells, both of which are located laterally on either side of the animals, are polarized posteriorly and anteriorly. This is followed by a second phase, in which the QL and QR cells migrate posteriorly and anteriorly, respectively, while undergoing the stereotyped cell divisions [34](Fig.2A). The first phase is controlled by at least three partially redundant genetic pathways, which are defined by genes encoding: (1) the transmembrane cell adhesion molecule UNC-40/DCC (Deleted in Colorectal Cancer)[41], (2) the transmembrane protein MIG-21 and the conserved, putative C-mannosyl transferase DPY-19/DPY19L [41–43], and (3) the cadherin proteins CDH-3 and CDH-4 [44–46]. Mutations in any of these genes result in defects in the initial polarization and, consequently, migration. Because *catp-8* mutants affected primarily QL migration, we focused our analysis on PVM and PQR migrations. We found that *unc-40; catp-8* double mutants exhibited a more severe phenotype than either single mutant alone (Fig.6B,D). Similar results were found in *mig-21; catp-8* double mutant animals, which also showed a more severe phenotype compared to the single mutants (Fig.6B,D). We could not test a double mutant with *cdh-*4, because the *catp-8; cdh-4* double mutant appeared lethal (data not shown). Therefore, we conclude that *catp-8* likely functions in redundant genetic pathways with *unc-40/DCC* and *mig-21*, respectively, to affect different neuronal migrations.

The *mab-5/HOX-C* antennapedia-like homeobox transcription factor is necessary and sufficient in QL cells after the initial polarization to mediate the second phase of their posterior migration [47] and is activated in response to the *egl-20/Wnt* morphogen that is expressed in posterior tissues [36]. Similar to loss of *catp-8*, loss of *mab-5* results in QL cell migration defect, where QL cell descendants migrate in an anterior rather than posterior direction [47]. Because the *mab-5* mutant phenotype is essentially fully penetrant, we could not test for possible enhancement in a *mab-5; catp-8* double mutant. In contrast, in a *mab-5(gf)* gain of function allele *mab-5* is activated in both QL and QR cells. However, migration of the QL descendants is not affected in the *mab-5*(gf) gain of function mutants, whereas QR descendants fail to migrate in an anterior direction [47]. We therefore tested the epistatic relationship between the *mab-5(gf)* allele and the *catp-8* loss of function mutant. We found that the aberrant anterior migration of the QL descendant PVM in *catp-8* single mutants was completely suppressed in a *mab-5(gf); catp-8* double mutant, whereas the migration defects of QR descendants remained unaffected (Fig.6C,E,F). Thus, we conclude that *catp-8/P5A-ATPase* functions upstream of or in parallel to the *mab-5/HOX-C* homeobox transcription factor.

### A functional CATP-8/P5A ATPase fusion localizes to the endoplasmic reticulum

Previous work showed that the yeast Spf1 (and Arabidopsis MALE GAMETOGENESIS IMPAIRED ANTHERS (MIA) or PDR2) homologs are localized to the endoplasmic reticulum (ER)[19, 20, 48]. We therefore aimed to determine the subcellular localization of CATP-8 in *C. elegans.* To this end, we used the functional N-terminal fusion of tagRFP to CATP-8 and drove expression from a single copy insertion transgene under control of the heterologous *dpy-7p* promoter, which is specific for the epidermis [49]. We chose to use this single copy, functional transgene over the endogenously tagged *catp-8* locus to minimize interference with signal from other tissues (cf. Fig.3). This epidermally-expressed transgene (*dzSi3*), which also displayed perinuclear staining (and rescued PVD patterning defects) was crossed with a panel of organellar reporters - for (1) the cis/medial Golgi, (2) ER, (3) early endosomes, (4) late endosomes and lysosomes, (5) autophagosomes, and (6) lysosomes. We found strong colocalization with a marker of the ER, but not with other vesicular markers (Fig.7). These findings suggest that in metazoans CATP-8/P5A-ATPase is localized to the ER similarly to plants and unicellular eukaryotes. We next tested whether the gross morphology of these cellular compartments was defective in *catp-8* mutant animals. Where tested, we found no apparent defects in the perinuclear localization, patterns or intensity of the tagRFP::CATP-8 transgene reporter in the epidermis (Fig.S6). Taken together, our findings suggest that the CATP-8/P5A-ATPase serves a function in the ER, which may be conserved from unicellular eukaryotes and plant to animals.

**Figure 7.**
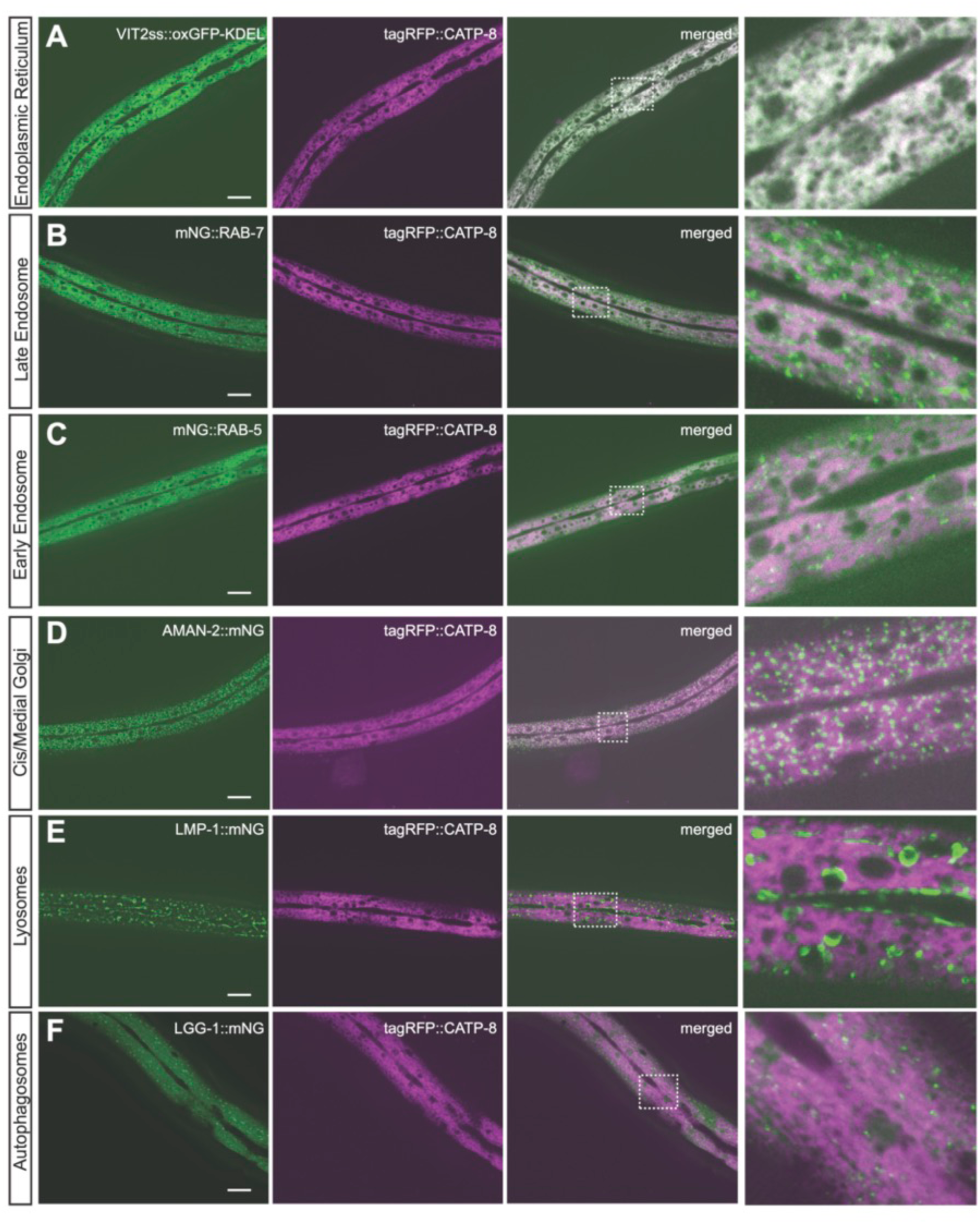
CATP-8/P5A-ATPase localizes primarily to the endoplasmic reticulum. A. - G. Confocal images of animals expressing a functional *tagRFP::CATP-8* fusion under control of the epidermal specific promoter *dpy-7* from a single copy transgene, in combination with single copy vesicular markers for the endoplasmic reticulum (A, *pwSi82[Phyp-7::VIT2ss::oxGFP-KDEL]*), late endosomes and lysosomes (B, *pwSi140[Phyp7::mNG::RAB-7;HYG-R]*), early endosomes (C, *pwSi145[Phyp7::mNG::RAB-5;HYG-R]*), cis/medial Golgi (D, *pwSi202[Phyp7::AMAN-2::mNG;HYG-R]*), lysosomes (E, *pwSi205[Phyp7::LMP-1::mNG;HYG-R]*), and autophagosomes (F, *pwSi144[Phyp7::mNG::LGG-1;HYG-R]*. The right column represents magnifications of the insets indicated.

### CATP-8/P5A ATPase is required for localization of some cell surface transmembrane molecules

The functions of *catp-8* in PVD neurons and the localization to the ER prompted us to test whether the localization or trafficking of two transmembrane molecules, which are expected to transit the ER, is affected in *catp-8* mutants. Specifically, we analyzed the localization of reporters for the single transmembrane receptor DMA-1/LRR-TM (DMA-1::GFP) [50] and the four transmembrane, claudin-like molecule HPO-30/Claudin (HPO-30::tagBFP) [51]. We found that overall DMA-1::GFP expression was noticeably dimmer in *catp-8* mutants (Fig.8A,B). Specifically, fluorescence in the cell body and the primary dendrite was strongly reduced in *catp-8* versus control animals (Fig.8C,D). Interestingly, the ratio of fluorescence in the cell body versus the primary dendrite was significantly increased (Fig.8E). These observations could be a reflection of reduced trafficking towards the periphery or, possibly, reduced amounts of DMA-1::GFP/vesicle. To distinguish between these possibilities, we quantified the number of puncta, previously suggested to represent the vesicular fraction of DMA-1::GFP [52, 53], in primary and tertiary dendrites of *catp-8* and control animals. We found the number of puncta of DMA-1::GFP in neither primary nor tertiary dendritic branches significantly different in *catp-8* mutant versus control animals. We therefore propose that *catp-8* does not serve a function in vesicle trafficking. Instead, *catp-8* may function to facilitate membrane localization of sufficient amounts of DMA-1/LRR-TM. Alternatively, but not mutually exclusive, *catp-8* could serve a role in loading DMA-1::GFP into vesicles.

**Figure 8.**
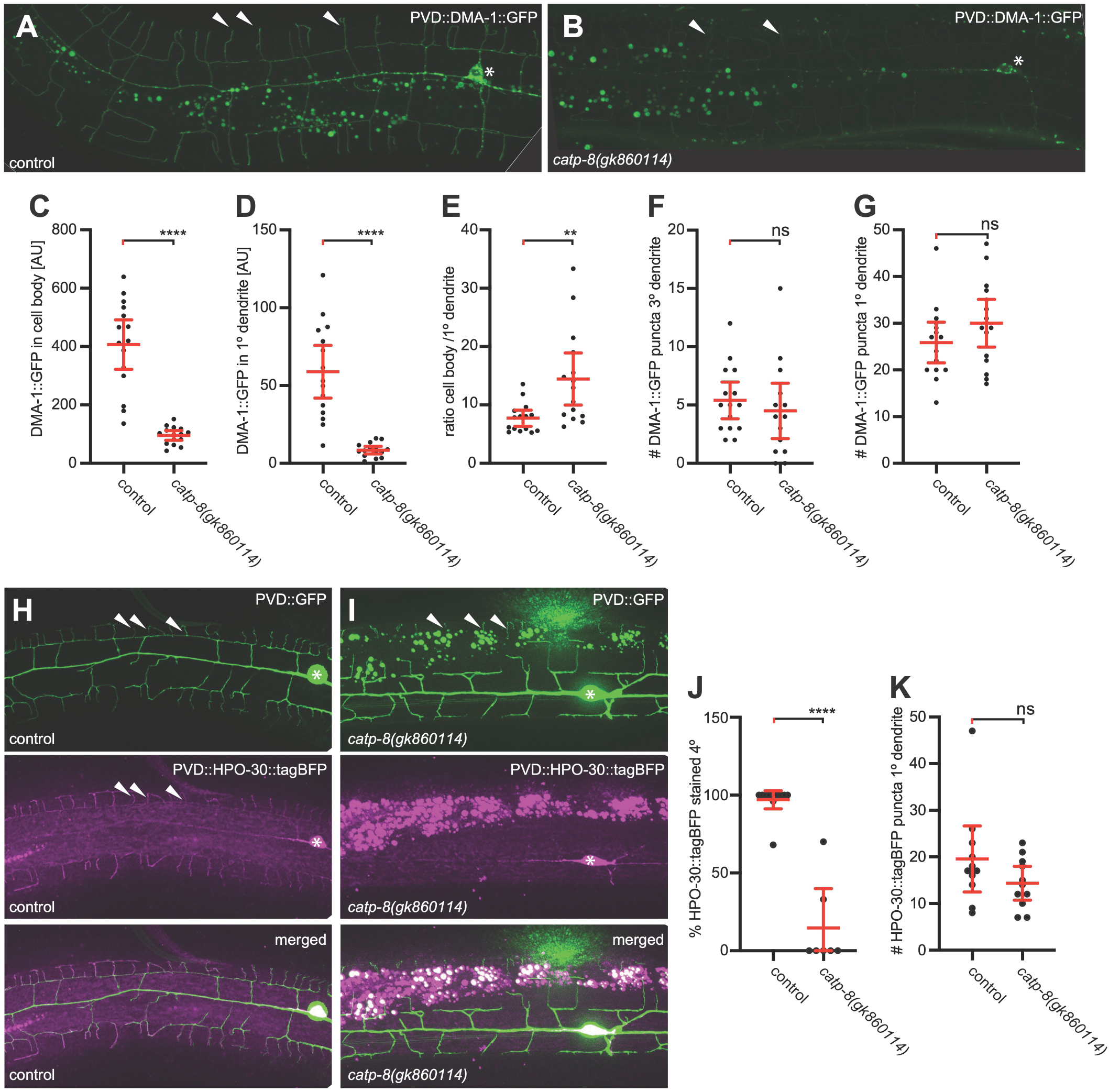
The CATP-8/P5A-ATPase is required for correct localization of transmembrane proteins in PVD dendrites. A. - B. Maximum z-projection of confocal images of DMA-1::GFP expressed in PVD (*qyIs369*) in a wildtype or *catp-8* mutant background, showing a decrease in overall fluorescence intensity of DMA-1::GFP in mutant animals. C. - E. Quantification of DMA-1::GFP signal in cell body (C), and on the primary dendrite 100 micron anterior from the cell body (D) in control and *catp-8* animals. The signal ratio of cell body to primary dendrite is presented in (E). Data are represented as the mean ± 95% confidence interval. *\*\* P<0.01, **** P<0.0001*, ns, not significant, Mann-Whitney test. n = 14 animals per genotype. F. - G. Quantification of DMA-1::GFP puncta either in the primary (F) or on the tertiary (G) dendrite 100µm anterior from the cell body in the genotypes indicated. ns, not significant, Mann-Whitney test. n = 14 animals per genotype. H. - I. Maximum z-projection of apotome images of HPO-30::tagBFP (magenta) expressed in PVD and PVD reporter *wdIs52* (green) in control (H) or *catp-8* mutant animals (I). Note, that as previously reported [51], overexpression of HPO-30::tagBFP in PVD dendrites results in a reduced number of quaternary dendrites, possibly due to downregulation of DMA-1/LRR-TM. J. Percentage of HPO-30::tagBFP stained quaternary dendrites 100 µm anterior to the cell body in control or *catp-8* mutant animals, presented as mean ± 95% confidence interval. *\*\*\*\* P<0.0001*, Mann-Whitney test. n = 10 animals per genotype. K. Number of HPO-30::tagBFP puncta in the primary dendrite 100 µm anterior to the cell body in control or *catp-8* mutant animals, presented as mean ± 95% confidence interval. ns not significant, Mann-Whitney test. n = 10 animals per genotype.

As previously reported, HPO-30::tagBFP localizes primarily to primary and quaternary dendritic branches [51]. We found that HPO-30::tagBFP fluorescence in distal dendrites appeared reduced in *catp-8* mutant animals (Fig.8H,I). Although HPO-30::tagBFP expression in the cell body did not seem visibly different, HPO-30::tagRFP fluorescence in quaternary dendrite branches was strongly reduced in *catp-8* mutant versus control animals (Fig.8J). Yet, the number of HPO-30::tagRFP puncta in the PVD primary branch remained unaffected, implying again that trafficking of HPO-30::tagBFP does not require *catp-8* function (Fig.8K).

Because epidermal expression of *catp-8/P5A-ATPase* could rescue PVD defects in *catp-8* mutant animals, we tested, whether another conserved transmembrane cell adhesion molecule SAX-7/L1CAM, which functions in the epidermis to shape PVD dendrites, is affected in *catp-8* mutant animals. We found that SAX-7::GFP localization in *catp-8* mutants was not visibly changed compared to control animals, either in neurons or the epidermis (Fig.S7A). We therefore conclude, *catp-8/P5A-ATPase* function is required for the regulation of some but not all transmembrane proteins.

## DISCUSSION

In this study we describe multiple functions of the *catp-8/P5A-ATPase* in metazoan development. Specifically, we show that *catp-8* is required for different aspects of nervous system development, including axonal guidance, dendrite patterning and neuronal migrations. A reporter for CATP-8/P5A-ATPase is expressed in the ER of most if not all tissues, and can function both cell-autonomously and non-autonomously during neuronal development. Genetically, *catp-8/P5A* functions in multiple pathways, including the conserved Menorin pathway during dendrite development and Wnt signaling during neuronal migrations. We find a substantial degree of genetic redundancy with different developmental pathways, suggesting that *catp-8/P5A-ATPase* may serve diverse, yet specific functions in different cellular contexts Lastly, *catp-8/P5A-ATPase* is necessary to correctly localize reporters for some but not all transmembrane reporters.

### Functional redundancy of catp-8/P5A-ATPase in neuronal patterning

Not all neurons are affected by loss of *catp-8* function. For example, the PVD somatosensory neurons require *catp-8* for the elaboration of dendritic trees, but the dendrites of amphid sensory neurons seem unaffected. Similarly, axonal guidance or extension of HSN and D-type motor neurons requires *catp-8* function, whereas guidance of PVQ axons or DA/DB motor neurons appears independent of *catp-8*, respectively. This observation is remarkable because PVQ and HSN (but also D-type and DA/DB motor neurons to some extent) make similar navigational choices, are part of the same fascicles and depend on similar axon guidance pathways, such as, Netrin signaling or Slit/Robo signaling [39, 54, 55]. It should be noted that Wnt signaling is also required for correct axon guidance of both HSN and PVQ axons at the midline [39, 56] and, at least for the latter cellular context, has been shown to function in parallel with Netrin signaling [56]. A possible explanation is that different genetic redundancies for *catp-8* exist in different cells; in other words, in some cells, *catp-8* may serve unique roles for which no functional alternative to *catp-8* exists whereas in other cells *catp-8* has functions that can be partially served by other genes. Alternatively, the molecules that are required for development of a given cell may be distinct in different neurons, such that some neurons require more or less proteins that depend on *catp-8* to function.

Transgenic rescue experiments suggest both cell-autonomous and non-autonomous functions for *catp-8.* For example, PVD can develop normally with *catp-8/P5A-ATPase* function present in either neurons or muscle. This finding is in contrast to findings by Feng et al. who could rescue *catp-8* defects by transgenic expression of *catp-8* in neurons, but not in muscle [23]. One possible explanation for this discrepancy is different levels in transgene expression (we use a single copy transgenes for epidermal expression whereas Feng et al. utilized multi copy transgenes [23]). This hypothesis is supported by the fact that the Menorin pathway is known to be dosage sensitive, and can be both positively and negatively regulated [57, 58].

In contrast to this tissue redundancy, the migration defects of Q cell descendants are rescued only by non-autonomous expression of *catp-8* in muscle, but not by expression in neurons or other tissues. These results strongly argue for a non-autonomous function of *catp-8/P5A-ATPase* in muscle during migration. However, no transmembrane or secreted molecules are known to function from muscle in Q cell migration, although muscle is clearly important, as the RNA-Binding Protein ETR-1/CELF1 can function in muscle cells [59]. It is therefore tempting to speculate that the EGl-20/Wnt ligand could be regulated by *catp-8/P5A-ATPase.* First, we found, that during HSN migration (but possibly also Q cell migration), Wnt signaling functions in concert with the *catp-8/P5A-ATPase.* Second, EGL-20/Wnt has been shown to be expressed from posterior body wall muscle among other posterior tissues [37, 38]. Lastly, the cellular migrations that are affected in *catp-8/P5A-ATPase* mutant animals (PVM, PQR, HSN) are all dependent on *egl-20/Wnt* whereas cell migrations that are independent on *catp-8/P5A-ATPase* (ALM, coelomocytes) are also independent of *egl-20/Wnt* [13]. Further experiments are needed to definitively establish whether EGL-20/Wnt secretion or localization is dependent on *catp-8/P5A-ATPase* function.

### Substrates of the CATP-8/P5A-ATPase

Previous studies in yeast and plants suggested various roles for P5A-ATPases, such as functions in phosphate homeostasis and male gametogenesis [18–20], as well as phospholipid and sterol homeostasis [21] and targeting of mitochondrial outer membrane proteins [22]. Therefore, which functions P5A ATPases may serve in metazoans remains a major question. Here, we provide genetic evidence that *catp-8/P5A-ATPase* is required for the localization of sufficient amounts of the DMA-1/LRR-TM receptor in PVD during dendrite morphogenesis. Similarly, the four transmembrane protein HPO-30/Claudin, which also functions in PVD neurons during dendrite patterning [60] requires *catp-8* to be correctly localized to distal, higher order dendrites. Importantly, the number of vesicles containing the respective reporters and, which are presumed to represent endocytic/transport vesicles [52, 53] along the dendritic branches, remain unaffected. Intriguingly, however, localization of the conserved SAX-7/L1CAM cell adhesion molecule in either neurons or the epidermis is not obviously affected. Therefore, *catp-8* may be involved in the insertion/localization of certain transmembrane proteins but not others to the plasma membrane. These findings are consistent with another recent study showing similar effects on DMA-1::GFP [23, 24], although it was also reported by Qin et al. that the levels of a HPO-30 reporter remained unchanged [24]. A possible explanation for this discrepancy is that we measured fluorescence in distal dendrites rather than the cell body [24]. The findings of a role for the *catp-8/P5A ATPase* in the localization of some cell surface proteins is seemingly at odds with two other studies in yeast and *C. elegans* suggesting that P5A-ATPases primarily maintain ER homeostasis by functioning as dislocases to remove mislocalized mitochondrial proteins [24, 25]. However, several arguments can be made that P5A-ATPases serve additional functions beyond removing mislocalized mitochondrial proteins. First, genetic manipulations in *C. elegans* that suppress ER defects in PVD as a result of the mislocalized DRP-1/Drp1 mitochondrial fission protein to the ER in *catp-8* mutants, fail to restore the correct dendritic morphology [24] suggesting that *catp-8/P5A-ATPase* serves additional functions beyond maintaining ER integrity to mediate PVD morphogenesis. Second, proteomic studies in HeLa cells found not only the levels of mitochondrial, but also other transmembrane and secreted proteins changed in ATP13A1/P5A-ATPAse mutant cells. For example, a number of secreted growth factors, including the platelet derived growth factor D (PDGFD) or transforming growth factor beta (TGFB) as well as axon guidance molecules such ephrin A5 (EFNA5), to name but a few, were significantly reduced in abundance upon loss of ATP13A1 catalytic activity [25]. In the context of these studies, our findings argue for a more diverse, yet likely specific role for *catp-8* in the localization of certain transmembrane proteins and regulation of secreted proteins. Further experiments will be required to delineate the full spectrum of genetic and biochemical functions of P5A-type ATPases.

## MATERIALS AND METHODS

### *C. elegans* strains and genetics

All *C. elegans* strains were grown on nematode growth medium plates with *E. coli* (OP50) as a food source as described [61], usually at 20°C unless otherwise specified. Strains used in this work can be found in Table S1. The *catp-8* alleles *dz224* and *gk860114* were backcrossed four and three times, respectively, prior to further analysis.

### Molecular cloning of *catp-8(dz224)*

Using a combination of Hawaiian SNP mapping and sequencing [26] the allele *dz224* was localized to a region between 8Mb and 13Mb on LG IV, which was further refined by deficiency mapping to between 10.2Mb and 12Mb (Fig.S1). This 1.8Mb interval comprised 7 non-synonymous missense mutations and one non-sense mutation in *catp-8*, which was confirmed by Sanger sequencing: an A to a T transversion at position 11,479,010 on LG IV (WB270), resulting in an ochre premature stop codon after Y55 in *C10C6.6* (*catp-8*). We found that *catp-8(gk860411)* failed to complement dz224, further suggesting that both alleles are affecting the same gene.

### Molecular biology and transgenesis

Cloning of all constructs were carried out using standard molecular biology methods. A list of primers used can be found in Table S2. For a list of plasmids used and detailed descriptions of constructs see Table S3 and S4. All extrachromosomal arrays were generated by injecting a mixture of the desired plasmids with *pBluscript* to a final concentration of 100 ng/μl of DNA. A detailed list of all the extrachromosomal transgenic lines generated can be found in Table S5.

For CRISPR/Cas9 mediated genome editing, appropriate guide RNA sites were selected using IDT Cas9 crRNA design tool and, oligos in the general form of TAATACGACTCACTATA(gRNA)GTTTTAGAGCTAGAAATAGCAAG were ordered, where (gRNA) is the 20nt of the guide RNA sequence before the PAM motif, optimized for T7 promoter transcription. These oligos were used in PCR reactions as a forward primer in conjunction with the universal reverse primer AAAAGCACCGACTCGGTG (oLT337) to generate sgRNA transcription templates from pDD162 (containing the sequence of tracrRNA). sgRNA was then transcribed from this template using HiScribe T7 *in-vitro* transcription kit (NEB) and, purified with Monarch RNA cleanup kit (NEB). The resulting sgRNA was used in an injection mix at 20 ng/µl per sgRNA, together with 250 ng/µl of Alt-R Cas9 endonuclease (IDT) and repair templates (100 ng/µl for single-strand oligos, 200 ng/µl for PCR products or plasmids). Injection and CRISPR efficiency were monitored through Co-CRISPR strategy with *dpy-10(cn64)* or *unc-58(e665)* conversions, or through *rol-6(su1006)* co-injection markers [62]. A list of all genome-edited strains and detailed methodology can be found in Table S6.

### Scoring of migration phenotypes

All worms were immobilized using 1 mM levamisole and mounted on 4% agarose pads for phenotypic analysis.

AQR/PQR migrations were scored in *wdIs52 (Is[F49H12.4p::GFP])* in L2 to young adults with a Plan-Apochromat 10x/0.25 objective on a Zeiss Axioimager Z1. Cells were considered in the head if anterior to the intestine, in the tail if posterior to the intestine, and in the mid body otherwise. AVM/PVM migrations were scored in *zdIs5 (Is[mec-4p::GFP])* in L4 to young adults with a Plan-Apochromat 10x/0.25 objective on a Zeiss Axioimager Z1. Cells were scored based on their relative position (anterior or posterior) to the vulva. HSNs migrations were scored in *dzIs75 (Is[kal-1p9::GFP])* in adult worms with a Plan-Apochromat 16x/0.5 objective on a Zeiss Axioimager Z1. Cells were considered to be at the vulva if the cell bodies were positioned within less than or equal to an egg’s length from the vulva. ALM was scored in *zdIs5 (Is[mec-4p::GFP])* in synchronized L1 animals. Animals are imaged using a Plan-Apochromat 10x/0.25 objective on a Zeiss Axioimager Z1 Apotome. Using ImageJ, the distance of the more anterior cell body from the nose was measured and divided by the total length of the animal. Coelomocytes were scored in *dzEx2140* at the L4 to young adult stage with a Plan-Apochromat 16x/0.5 objective on a Zeiss Axioimager Z1. Coelomocytes were considered to be in the head region if in the anterior quarter of the worm.

### Scoring of axon pathfinding HSN axon pathfinding

HSN axon guidance at the ventral midline was scored in *dzIs75* at the L4 to adult worms stage with a Plan-Apochromat 40x/1.3 objective on a Zeiss Axioimager Z1. Crossover defects of axons were scored if any portion of the axons of HSNL and HSNR were touching. DD/VD neurons were scored in *juIs76* at the L4 to adult worms stage with a Plan-Apochromat 40x/1.3 objective on a Zeiss Axioimager Z1. Axon guidance was categorized into two measurements, L/R choice and gaps in the dorsal or ventral cord. For L/R choice, any worm with 1 or more axons found on the opposite side of the stereotyped location was counted as having an L/R choice defect. Additionally, any worm with a visible discontinuation of the dorsal and/or ventral cord between any two neurons (gaps) was considered to have a dorsal and/or ventral cord defect respectively. DA/DB motor neurons were scored in *evIs82b* at the L4 to adult stage with a Plan-Apochromat 40x/1.3 objective on a Zeiss Axioimager. Axon guidance was categorized into two measurements, L/R choice and gaps in the dorsal or ventral cord. For L/R choice, any worm with 1 or more axons found on the opposite side of the stereotyped location was counted as having an L/R choice defect. Any worm with a visible discontinuation of the dorsal and/or ventral cord between any two neurons (gaps) was considered to have a dorsal or ventral cord defect, respectively. AIY axonal morphology was scored with *mgIs32* at the L4 to adult stage with a Plan-Apochromat 16x/0.5 objective on a Zeiss Axioimager Z1. Cells were considered to be overbranching if an additional protrusion off of the cell body at least the length of the cell body was present. PVQ axonal guidance was scored in *oyIs14* at the L4 to adult stage with a Plan-Apochromat 16x/0.5 objective on a Zeiss Axioimager Z1. Crossover defects were counted if any portion of the axons of PVQL and PVQR anterior to the vulva were not visibly separable. For DiI staining**, w**ell-fed adult worms were washed from plates with 1 mL M9, pelleted, resuspendend in 1 mL M9 before 5 µL of DiI stock solution (2 mg/mL DiI (Molecular Probes, #D282) in dimethyl formamide) was added. Animals were incubated with light shaking for 3 hours, spun and washed with M9 two times, followed by transfer to fresh plates. After 30 minutes recovery, animals were immobilized in 1 mM levamisole and neuronal processes were assessed for defects and imaged with a Plan-Apochromat 40x/1.3 or 63x/1.4 objective on a Zeiss Axioimager Z1 Apotome.

### Fluorescent microscopy and quantification

#### PVD morphometric analysis

Fluorescent images were captured in live *C. elegans* at the L4 larval stage using a Plan-Fluor Nikon 40x/1.3x on a Nikon CSU-W1 Spinning Disk Confocal. At least 15 adult animals were scored per genotype. Optical sections were collected and maximum intensity projections adjusted for optimal contrast to resolve detailed morphology were used for further analysis. For tracing, the 100μm section of the primary branch anterior to the cell body was used for morphometric analyses using the NeuronJ plugin of the FIJI software. Branches were defined as follows: 2° dendritic branches as any neurite branching out of the primary dendrite; 3° branches as neurites branching at the end of a 2° branch in proximity to the tertiary line along the border of outer body wall muscles and the lateral epidermis; 4° dendritic branches as those originating from 3° dendritic branches and extending towards the dorsal or ventral nerve cords, respectively.

#### DMA-1::GFP quantifications

All DMA-1::GFP fluorescent images were captured in strains containing *qyIs369* at the L4 larval stage using a Plan-Fluor Nikon 60x/1.4x objective and GFP filter setting on a Nikon CSU-W1 Spinning Disk Confocal with identical imaging parameters and appropriate z-stack. Fluorescence measurements of the z-projected image were subsequently performed in ImageJ as follows. For Intensity of GFP in the PVD cell body an outline of the cell body was drawn as Region of Interest (ROI) and the average signal in the ROI was quantified. An ROI of the same shape was placed outside of the cell body to measure the background signal, which was subtracted from the fluorescent signal. For the intensity of GFP in the PVD primary dendrite, the primary dendrite was traced for 100 µm anterior to the cell body, and an ROI was defined as 10 pixels (= 1.6 µm) thick along the traced line, where the average signal in the ROI was quantified. A ROI of the same size was placed next to the dendrite to measure the background signal, which was subtracted from the fluorescent signal. The number of puncta within 100 µm of the primary dendrite anterior to the cell body were counted manually.

#### HPO-30::tagBFP quantification

All HPO-30::tagBFP fluorescent images were captured in strains containing *dzEx2140* and *wdIs52* at the L4 larval stage using a 40x/1.3 objective on a Zeiss Axioimager Z1 Apotome with identical imaging parameters for tagBFP and GFP filter setting, and appropriate z-stacks. Quantification of the z-projected images were subsequently performed in ImageJ as follows

For the percentage of quaternary dendrites containing HPO-30::tagBFP, the number of quaternary branches with visible tagBFP signal within 100 µm anterior to the cell body were counted in the tagBFP channel, then divided by the number of quaternary dendrites as imaged in the GFP channel. For the number of HPO-30::tagBFP puncta, quantification was performed by manually counting the puncta within 100 µm anterior to the cell body.

#### SAX-7::GFP imaging

Fluorescent images for the strains containing the *ddIs290* fosmid based SAX-7::GFP reporter were captured at the late L4 larval stage using a Plan-Apochromat 40x/1.3 objective on a Zeiss Axioimager Z1 Apotome at an exposure time of 151 ms, followed by appropriate z-stacks. Neuronal and hypodermal patterning were assessed, and the PVD phenotype was observed in a *dzIs53* background.

#### Colocalization assay of subcellular compartment markers

L4 animals carrying *dzSi3* and single-copy insertions of GFP subcellular compartment markers were imaged using Plan-Fluor Nikon 60x/1.4x on a Nikon CSU-W1 Spinning Disk.

### Statistical Analysis

All statistical analyses were performed using the Prism 9 Statistical Software suite (GraphPad). All pairwise migration phenotypes are analyzed with Fisher exact test when applicable, or Chi-squared test as indicated. All comparisons of averages/proportions are analyzed with one-way ANOVA with Tukey correction, the Kruskal-Wallis test, the Mann-Whitney test, or the Z-test as applicable.

## ACKNOWLEDGMENTS

We thank S. Emmons, A. Meléndez, Y. Salzberg, and members of the Bülow laboratory for comments on the manuscript and for helpful discussions during the course of this project. We thank S. Cook for the initial observation of PQR cell migration defects, R. Poole for help with the analysis of whole genome sequencing data and, Yuji Kohara for cDNA clones; some strains were provided by the CGC, which is funded by NIH Office of Research Infrastructure Programs (P40 OD010440). We acknowledge help with imaging from the Analytical Imaging Facility at Albert Einstein College of Medicine, which is supported in part by the NCI Cancer center grant (P30CA013330). In addition, this work was supported by grants from the NIH (R01NS096672 and R21NS111145 to HEB, F31NS111939 to MT, F31NS100370 to MR, T32GM007491 to CJS, P30HD071593 to Albert Einstein College of Medicine, and R01GM135326, R01GM067237, and R01AG047101 to BDG). LTHT was the recipient of a Croucher Foundation Research Fellowship and JF of a Tishman Scholarship from the Dominick P. Purpura Department of Neuroscience.

## Supplementary Figures

**Figure S1 Mapping and auxiliary genetics**

A. Frequency plot of N2 SNPs from whole genome sequencing results as per Hawaiian SNP mapping [26], indicating *dz224* lesions reside between 6 Mbp and 13 Mbp on linkage group (LG) IV. Deficiencies covering the region indicated below (green, complementation of *dz224*; red non-complementation of *dz224*), demonstrating that *dz224* lesions lies between 10,202,993, and 12,011,007 bp. Table below listing lesions detected within this range, with *catp-8* containing the only stop-gain mutation.

B. - E. Quantification of the number of non-ectopic secondary branches (E), aggregate length of secondary (F), tertiary (G), and quaternary (H) dendrite branches 100µm anterior to the PVD cell body in the genotypes indicated. Data are represented as the mean ± 95% confidence interval. Statistical significance was calculated using one-sided ANOVA with Tukey’s multiple comparison test. *, *P* ≤ 0.05; ***, *P* ≤ 0.001; ****, *P* ≤ 0.0001; ns, not significant.

F. Complementation assay tallying percentage of animals with PVD defects, showing non-complementation between *dz224* and *catp-8(gk860114),* and recessiveness of *dz224* and *catp-8(gk860114)*.

G. Fraction of *catp-8* homozygous animals presenting PQR migration defects from homozygous or heterozygous *catp-8* mutant parent. PQR migration is completely normal in F1s of heterozygous animals, showing maternal contribution of *catp-8* in PQR migration. Homozygosity of animals was determined by the PVD phenotype, which is not maternally rescued.

H. Quantification of number of quaternary branches, secondary branches, and the ratio of quaternary to secondary branches 100µm anterior to the PVD cell body in *catp-8(gk860114)* and *catp-8(gk860114)/mDf7* transheterozygous animals. No enhancement was observed in the transheterozygote, indicating *gk860114* is a genetic null allele. Data are represented as the mean ± 95% confidence interval. ns, not significant, Kruskal-Wallis test with Dunn’s multiple comparisons test. n = 14 animals per genotype.

I. L1 body length of wildtype and *catp-8(gk860114)* measured and presented as mean ± 95% confidence interval. ns not significant. Statistical significance was calculated with an Unpaired t-test.

**Figure S2 Additional neuronal phenotypes in *catp-8* mutant animals**

A. - B. Maximum z-projection confocal images DA/DB cholinergic motorneurons (visualized by *evIs82b(Is[unc-129p::GFP]*) of wildtype (A) and *catp-8* (B) animals. No difference between the genotypes were observed. Lateral views, ventral up, anterior to the left.

C. - D. Maximum z-projection confocal images D-type GABAergic motorneurons (visualized by *juIs76(Is[unc-25p::GFP]*) of wildtype (C) and *catp-8* (D) animals. Occasional gaps caused by under-extension of neurite can be observed in *catp-8* mutant animals. Lateral views, dorsal up, anterior to the left.

E. - F. Maximum z-projection confocal images of AIY interneurons (visualized by *mgIs32(Is[ttx-3p::GFP]*) in wildtype (E) and *catp-8* (F) animals. No difference between the genotypes were observed. Lateral views, dorsal up, anterior to the left.

G. - J. Maximum z-projection apotome images of DiI stained phasmid sensory neurons in wildtype (G) and *catp-8* (H) animals, and amphid sensory neurons in wildtype (I) and *catp-8* (K) animals. No difference between the genotypes were observed. Ventral views, anterior to the left. Lateral views, dorsal up, anterior to the left.

K. - N. Maximum z-projection confocal images of PVQ interneurons (visualized by *oyIs14(Is[sra-6p::GFP])* (K.,L.) and HSN motorneurons (visualized by *dzIs75(Is[kal-9p::GFP]*) (M..N.) in wildtype and *catp-8* animals. No defects were observed in PVQ axons at the midline, but HSN neurons showed a significant number of midline cross overs in *catp-8* mutant animals. Ventral views, anterior to the left.

**Figure S3 A *catp-8* transcriptional reporter is widely expressed**

A. - E. Maximum z-projection confocal images of animals at embryo (A), L2 (B), L3 (C-D) and L4 (E) stages of worms carrying *catp-8* promoter GFP fusion array *dzEx2101*. *dzEx2101* exhibits high degree of mosaicism, resulting in staining of different tissues across the population, including muscle, intestine, pharynx, and some neurons.

**Figure S4 Complete data of cell specific rescue experiments**

A. - C. Quantification of number of quaternary branches (A), secondary branches (B), and the ratio of quaternary to secondary branches (C) 100µm anterior to the PVD cell body in animals of all extrachromosomal rescue lines tested. Data are represented as the mean ± 95% confidence interval. *\*\*\*\* P<0.0001*, ns, not significant, Kruskal-Wallis test with Dunn’s multiple comparisons test. n = 14 animals per genotype.

D. - E. Percentage of animals with the indicated migration phenotype of PVM (D) and PQR (E) for all extrachromosomal rescue lines tested. **** *P < 0.0001*, ns not significant, Chi-squared test. n > 75 animals per genotype.

**Figure S5 Additional morphometric data of genetic double mutant analysis**

A. - C. Quantification of the aggregate length of secondary (A), tertiary (B), and quaternary (C) dendrite branches within 100µm anterior to the PVD cell body in animals of the indicated genotypes. Note that all alleles are molecular or genetic null alleles. Data are represented as the mean ± 95% confidence interval. *\* P<0.05, ** P<0.01, *** P<0.001, **** P < 0.0001* ns not significant; one-sided ANOVA with Tukey’s multiple comparison test. n = 15 animals per genotype.

**Figure S6 Different organelles are not visibly affected in *catp-8* mutants**

Confocal images of single copy insertion transgenes of epidermally expressed organellar reporters in *catp-8(gk860114)* mutant and wild type animals, for late endosomes (*pwSi140[Phyp-7::mNG::RAB-7; HYG-R]*), early endosomes (*pwSi145[Phyp-7::mNG::RAB-5; HYG-R]*), cis/medial Golgi (*pwSi202[Phyp-7::AMAN-2::mNG; HYG-R]*), and lysosomes (*pwSi205[Phyp-7::LMP-1::mNG; HYG-R]*).

**Figure S7 SAX-7::GFP localization is not affected and OE does not rescue PVD defects in *catp-8* mutants**

A. Fluorescent images of animals expressing a functional fosmid based SAX-7::GFP reporter (*ddIs290*) in control and *catp-8(gk860114)* mutant animals. White arrowheads denote neuronal staining, while open arrows indicate examples of epidermal staining at the lateral epidermal ridge.

B. Quantification of the percentage of animals with PVD 4° branching defects in indicated genotypes in a *ddIs290* background. Statistical comparisons were performed using the Z-test. Statistical significance is indicated (\*\*\*\**P<0.0005*). n = 15 animals per genotype.

